# Eliminating the Missing Cone Challenge through Innovative Approaches

**DOI:** 10.1101/2024.01.11.575283

**Authors:** Cody Gillman, Guanhong Bu, Emma Danelius, Johan Hattne, Brent Nannenga, Tamir Gonen

## Abstract

Microcrystal electron diffraction (MicroED) has emerged as a powerful technique for unraveling molecular structures from microcrystals too small for X-ray diffraction. However, a significant hurdle arises with plate-like crystals that consistently orient themselves flat on the electron microscopy grid. If, as is typically the case, the normal of the plate correlates with the axes of the crystal lattice, the crystal orientations accessible for measurement are restricted because the grid cannot be arbitrarily rotated. This limits the information that can be acquired, resulting in a missing cone of information. We recently introduced a novel crystallization strategy called suspended drop crystallization and proposed that this method could effectively address the challenge of preferred crystal orientation. Here we demonstrate the success of the suspended drop crystallization approach in eliminating the missing cone in two samples that crystallize as thin plates: bovine liver catalase and the COVID-19 main protease (Mpro). This innovative solution proves indispensable for crystals exhibiting preferred orientations, unlocking new possibilities for structure determination by MicroED.

## 1. Introduction

Microcrystal electron diffraction (MicroED) is a cryogenic electron microscopy (cryo-EM) method in which vanishingly small crystals are used for structure determination by electron diffraction (Shi *et al*., 2013). In MicroED, microcrystalline samples are prepared on cryo-EM grids and diffraction data are collected as a movie on a fast camera in a transmission electron microscope (TEM) while the sample is continuously rotated (Nannenga, Shi, Leslie *et al*., 2014). This setup allows the data to be processed using standard crystallographic software. The use of micro- and nanosized crystals provides new opportunities for determining previously unattainable structures, and MicroED has been applied to small molecules, natural products, materials, soluble- and membrane proteins, and is gaining popularity for various applications (Bruhn *et al*., 2021; Mu *et al*., 2021). For highly symmetric crystals, MicroED routinely yields complete data from single crystals, even for proteins as in the case for proteinase K (Martynowycz *et al*., 2023) and the adenosine A2a receptor (Martynowycz *et al*., 2023).

The experimental design of the sample stage in relation to the incoming electrons inherently limits the rotation range that can be used for data collection. At high tilt angles, the electron path through the sample is obstructed by grid bars or other crystals. Moreover, the grid holder itself blocks the electron beam at angles above∼70° so the maximum rotation range is∼140° not the ideal 180°. To address this challenge, numerous datasets must be collected, processed, and screened for merging statistics, aiming at completing reciprocal space (Griner *et al*., 2019; Gallagher-Jones *et al*., 2018 Danelius *et al*., 2023). Some examples include triclinic lysozyme, where data from 16 crystals were merged into a 88% complete dataset (Clabbers *et al*., 2022) and the human G-protein coupled receptor vasopressin, also crystallized in P1, where 14 datasets had to be merged to yield 75% completeness (Shiriaeva *et al*., 2023).

However, certain crystal morphologies pose specific challenges: when the crystals are flat and the orientation of the crystallographic axes follows the orientation of the plane, the crystal lattice will assume a preferred orientation because plate-like crystals tend to lie flat on the surface of the sample grid. In combination with the limited rotation range of the stage, this renders a systematic cone of reciprocal space inaccessible for data collection, irrespective of the number of datasets merged and the symmetry of the crystal (Glaeser & Downing, 1993; Stahlberg *et al*., 2015). This is the case for all examples of 2D crystals since they can only adopt a single orientation on the EM grid. For such samples, the maximum achievable completeness is∼86% even if the stage is rotated through the maximum possible range (Glaeser & Downing, 1993). Thus, the missing cone creates a substantial bottleneck in structure solution for crystals that adopt a preferred orientation. Missing wedges in reciprocal space result in artifacts in the density, such as elongated and/or missing or irrecoverable disjoint volumes (Stahlberg *et al*., 2015). Although common for micro- and nanosized crystals, this challenge is not unique to MicroED; but is well-known in transmission electron microscopy. For example, it manifests in cryotomography (Barth *et al*., 1988), where whole cells can align to the support grids. It should be noted that the incomplete crystal lattice sampling due to the missing cone can be eliminated if data from multiple crystals that are randomly oriented on the grid are combined.

Recently, we reported a new method for crystal growth called suspended drop crystallization (Gillman *et al*., 2023). With this approach crystals grow in suspension directly on a grid without any support film layered over the grid bars. Before data collection, the crystal and the surrounding material is FIB-milled to a thickness amenable to MicroED. Because no grid support film is present, we postulated that preferential orientation of crystals would be greatly reduced and that this approach could therefore eliminate the missing cone problem. Here, we demonstrate this by collecting complete datasets from two samples that are known for preferred orientation, namely bovine liver catalase, and the COVID-19 main protease (Mpro).

## 2. Methods and Materials

### 2.1 Protein expression and purification of Mpro

The gene encoding the full-length SARS-COV-2 Mpro was designed into the pGEX-6P-1 vector. The construct was fused with an N-terminal self-cleaving GST tag and a C-terminal His affinity tag downstream of a Precission protease cleavage site. *E. coli* strain Rosetta2 (DE3) was transformed with the expression plasmid for protein expression. An overnight culture was grown in terrific broth at 37 °C and used for inoculating 1 L of culture, which was incubated at 37 °C and 225 rpm. Protein expression was induced by 0.5 mM isopropyl β-d-1-thiogalactopyranoside (IPTG) when the OD_600_ reached 0.7. Expression was allowed for 16 h at 18 °C. The cell pellet was harvested at 8,000 g and kept at -20 °C until use. The frozen cell pellet was thawed at 4 °C in lysis buffer (50 mM HEPES-NaOH pH 7.5, 2 mM DTT, 2 mM EDTA, 2 mM EGTA, 0.1 mg/mL lysozyme and 0.1 mg/mL DNase I). Cells were lysed by sonication at 70% amplitude for 5 min. The cell debris and unbroken cells were removed by centrifuging at 12,000 g and 4 °C. The supernatant was directly loaded into a gravity column filled with 1 mL TALON resin (TaKaRa) pre-equilibrated with the binding buffer (20 mM HEPES-NaOH pH 7.5, 100 mM NaCl and 1 mM DTT). The column was washed with 25 mL wash buffer (20 mM HEPES-NaOH pH 7.5, 100 mM NaCl, 1 mM DTT and 10 mM imidazole), then washed with 10 mL Precission protease cleavage buffer (50 mM Tris-HCl pH 8.2, 150 mM NaCl and 1 mM DTT). Approximately 0.2 mg Precission protease in 3 mL cleavage buffer was directly added into the TALON resin slurry. The resin slurry was incubated at 4 °C for over 16 h with gentle shaking. The tag-cleaved Mpro was collected in fractions while flowing off the resin. The purified protein was concentrated to 0.6 mL using a 10 kD cut-off Amicon Ultra centrifugal filter (Millipore). The sample was then loaded into a Superdex 200 Increase 10/300 GL column (Cytiva) pre-equilibrated with the SEC buffer (20 mM Tris pH 7.8, 100 mM NaCl, 1 mM DTT and 1 mM EDTA). The collected Mpro fractions were pooled, concentrated using an Amicon Ultra centrifugal filter with 10 kD cut-off (Millipore), and stored at -80 °C until use.

### 2.2 Crystallization assays

Catalase was crystallized using procedures based on previously reported methods (Nannenga et al., 2014). First, a crystalline suspension of catalase from bovine liver (C100; Sigma-Aldrich) was centrifuged to pellet the crude catalase crystals. The supernatant was decanted and the pelleted crystals were washed in water and centrifuged again to recollect the catalase crystals. The supernatant was decanted and a solution of 1.7 M NaCl was added to solubilize the crystalline catalase. Solubilization was performed at room temperature for 1 h. The freshly solubilized catalase was dialyzed overnight at 4 °C against a solution of 50 mM sodium phosphate (pH 6.3) to allow recrystallization. The catalase crystals were collected and stored in tubes at 4 °C for an additional 24 h before being washed with water as described above. Catalase crystals were stored at 4 °C until used for MicroED sample preparation.

The Mpro fraction was thawed and the concentration was adjusted to 5 mg/mL. The mixture was diluted 1:1 with crystallization mother liquor (0.1 M MES pH 6.5, 20% PEG 3,350 and 5% DMSO). Crystals grew as plate clusters after 1 to 3 days when incubating at 20 °C using hanging vapor diffusion. Approximately 10 μL of the crystal drops were combined and used for preparation of a micro seeding stock. 1 μL of 2.5 mg/mL Mpro was mixed with 0.1 μL seeding stock and 1 μL of the crystallization condition. Monodisperse Mpro microcrystal plates were grown after 1 day when incubating at 20 °C using hanging vapor diffusion.

### 2.3 EM sample preparation

EM samples were prepared following previously published protocols (Martynowycz *et al*., 2021). Typically, carbon-coated (Quantifoil R 2/2) 200-mesh copper grids (Electron Microscopy Sciences) were negatively glow-discharged at 15 mA for 45 s using a PELCO easiGlow (Ted Pella). The grids were then loaded into a Leica EM GP2 plunge freezer (Leica Microsystems) set to 20 °C and 95% humidity. Using a micropipette, 2 μL of the crystal drop was transferred to the grid, which was then blotted for 20 s, and finally plunged into liquid ethane.

For the suspended-drop grids, the crystal slurry was added directly to a 150 mesh count support-free gold gilder grid (Ted Pella) that was clipped into an autogrid cartridge (Thermo-Fisher). To transfer a minimal amount of catalase sample to the grid, the sample was first aspirated into a pipet tip, which was gently dragged across the surface of the grid to deposit the sample. Sample transferred to the grid bars and was retained by surface tension. For MPro, the grids supporting the suspended crystal drops were blotted with filter paper from the edge for 2 s. The grids were finally plunged into liquid ethane and stored under liquid nitrogen until further use.

### 2.4 Milling lamellae of catalase suspended crystals

Focused ion beam (FIB) milling followed procedures similar to suspended drop crystallization (Gillman *et al*., 2023). The frozen sample containing suspended crystals was loaded into a Thermo-Fisher Helios Hydra plasma beam FIB (pFIB)/scanning electron microscope (SEM) operating at cryogenic temperature. The crystal drop was coated with platinum by beam-assisted GIS coating (argon beam at 4 nA, 5 kV) for 1 min to protect the sample from ion and electron beams. A whole-grid atlas of the drop was acquired with the SEM operating at an accelerating voltage of 0.5 kV and beam current of 13 pA using the latest version of MAPS software (Thermo-Fisher). The pFIB milling angle was set to 10° from the grid surface. For the first milling step, in the pFIB view, two 20×20 μm boxes (cleaning cross section milling pattern) separated by 5 μm were drawn and centered about the site of interest. Bulk material was removed using the xenon plasma beam set to a current of 4 nA for rough milling, which produced a 10-15 μm long lamella. The lamella trenches were inspected to ensure that the top and bottom of the lamella were exposed for subsequent MicroED data acquisition. The second lamella milling step used 1 nA of xenon plasma current to reach 3.5 μm thickness and 15 μm width. The third milling step was performed with a 0.3 nA xenon plasma current to narrow the lamella down to 2 μm thickness and 10 μm width. The fourth milling step was done at 0.1 nA to reach 1 μm thickness. Lastly, the milling ion current was set to 30 pA for final thickness milling and polishing. A 10 μm wide and roughly 300 nm thick lamella was generated through the drop of suspended crystals. The lamellae were visualized at 385 nm excitation wavelength with the iFLM on the Hydra dual-beam instrument.

### 2.5 Milling lamellae of Mpro preferred orientation crystals and suspended crystals

The vitrified grids were transferred into a Thermo-Fisher Aquilos FIB/SEM instrument operating at cryogenic temperature. Whole-grid atlases were recorded with the SEM at 5 kV and 1.6 pA using MAPS software. To protect the samples from the damaging ion and electron beams, the grids were coated with platinum by sputter coating (Martynowycz *et al*., 2019*b*). Individual crystals from blotted grids and suspended drops were identified and aligned to eucentric height using MAPS software. FIB-milling proceeded as described (Martynowycz *et al*., 2019*b*,*a*). A typical crystal identified on the blotted grids was tilted to a milling angle of 33°. Rough milling used an ion beam current of 300 pA to produce a 4 μm thick and 5 μm wide lamella. For fine milling, a FIB current of 100 pA was used to reduce the thickness of lamella to 1 μm. Polishing proceeded at an ion beam current of 50 pA until the thickness reached approximately 350 nm. The final lamella was approximately 3 μm wide. Milling suspended drop samples was similar to the previously mentioned steps with slight modifications: the sample was tilted to a milling angle of 13°, and the ion beam current for rough, fine, and final milling was set to 3 nA, 500 pA, and 100 pA, respectively.

### 2.6 MicroED data collection

The grids hosting the milled lamellae were rotated 90° and transferred to a cryogenically cooled Thermo-Fisher Scientific Titan Krios G3i TEM. The Krios was equipped with a field emission gun operating at an accelerating voltage of 300 kV, a Falcon 4i direct electron detector, and a Selectris energy filter (Thermo-Fisher). A low magnification atlas of the grid was acquired using EPU (Thermo-Fisher) to locate all milled lamellae. The stage was moved to the lamella position and while in the “View” settings (SA 3,600×) of SerialEM (Mastronarde, 2003), the eucentric height was set. The lamellae were manually scanned by sequentially evaluating electron diffraction (Figure 1) until a crystal site was located. In the “Record” settings, the following parameters were set for the electron beam in diffraction mode: a beam size of 20 μm in diameter, a spotsize of 11, and a C2 aperture of 50 μm. These settings resulted in an electron dose rate of approximately 0.0025 e-/(Å^2^·s). Electron-counted MicroED datasets were collected using a Falcon 4i with the “Record” mode of SerialEM following the protocol described in previously published work (Shiriaeva *et al*., 2023). An in-house developed script was applied to insert the Selectris energy filter and automatically collect continuous-rotation MicroED datasets for 420 s with a slit width of 20 eV and a selected area aperture diameter of∼2 μm. Autodoc and log files were generated after each data collection to provide metadata and parameters to use for data processing. For catalase, MicroED data were collected by continuously rotating the stage at 0.14 ° / s for 420 s, resulting in a rotation range of 60°. For Mpro, typical datasets from blotted crystals and suspended-drop grids were collected by continuously rotating the stage at 0.19 ° / s for 420 s, resulting in a rotation range of 80°.

**Figure 1.**
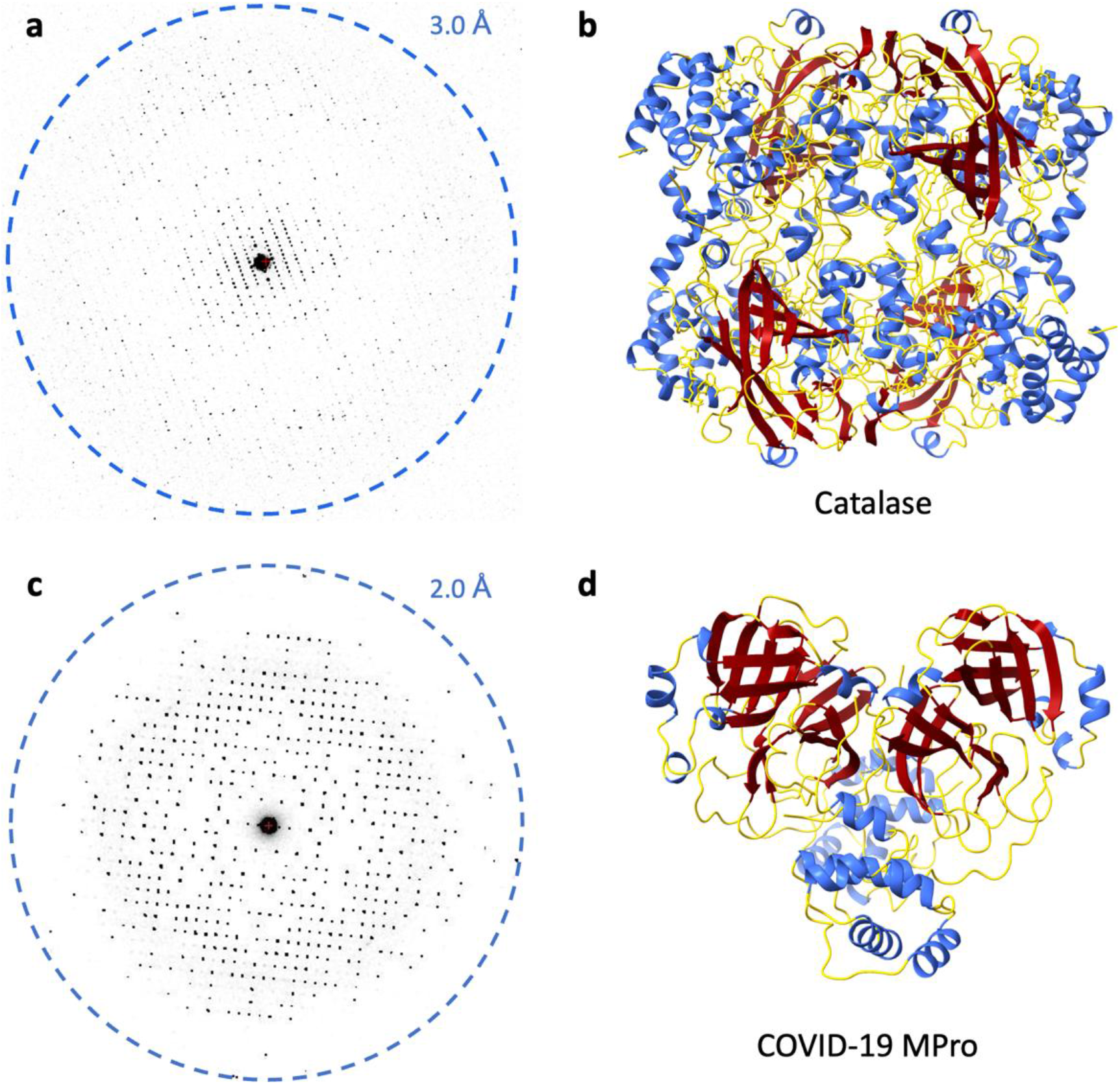
MicroED of catalase and Mpro. (a) Electron diffraction frame acquired from catalase. (b) Ribbon model of catalase. (c) Electron diffraction frame acquired from Mpro. (d) Ribbon model of Mpro.

### 2.7 MicroED data processing and structure determination

Movies were converted to SMV format using the latest version of MicroED tools (Martynowycz *et al*., 2019*a*; Hattne *et al*., 2015; Hattne *et al*., 2023). The converted MicroED datasets were processed with XDS (Kabsch, 2010*b*). The catalase diffraction dataset was indexed and integrated in space group *P* 2_1_ 2_1_ 2_1_ and unit cell dimensions 68.65, 173.30, 182.78 (a, b, c) (Å) and 90, 90, 90 (α, β, γ) (°), and the Mpro dataset was indexed and integrated in space group *C* 2 with the unit cell parameters of 115.54, 55.53, 45.37 (a, b, c) (Å), 90, 101.07, 90 (α, β, γ) (°). High quality datasets including the missing cone were scaled and merged in XSCALE (Kabsch, 2010*a*) and converted to MTZ format with supplemental 5% free R column in XDSCONV (Kabsch, 2010*b*). Phases for the MicroED reflections were determined by molecular replacement in MOLREP and Phaser (McCoy et al., 2007) using Protein Data Bank (PDB) 3J7B (Nannenga *et al*., 2014) as the search model for catalase, and PDB 7K3T (Andi *et al*., 2022) as the search model for MPro. The refinement was performed with REFMAC and phenix.refine (Afonine *et al*., 2012) using electron scattering factors without restoring missing reflections. Refinement was followed by manual curation in Coot (Emsley *et al*., 2010).

## 3. Results

### 3.1 Sample preparation and MicroED data collection

To collect data covering all of reciprocal space, sample preparations were screened for randomly oriented crystals. In addition, the samples were kept at minimal volume to reduce the FIB-milling time and thereby prevent damaging the crystals in the suspended drops. Initially, 0.3–0.5 μL of crystal slurry was pipetted onto the support-free grid and spread with a micro-brush. However, pipetting such small volumes gave inconsistent results using standard micropipettes. The second application method explored was preparing a droplet of the crystal slurry on a siliconized coverslip and gently touching a support-free grid to the surface of the crystal slurry, which transferred a fraction of the drop to the grid bars. Although this improved the consistency of sample preparation, spreading the drop with a micro-brush was still required. It was finally discovered that gently touching a pipet tip filled with the crystal slurry to the grid bars allows a minimal volume of the slurry to transfer by capillary motion. This method was highly reproducible and efficient leading to very small volume. We further evaluated careful blotting from the side with filter paper to reduce the volume of the sample, which also proved to be successful. The thickness of the samples from both methods was estimated to be similar to the height of the gold grid bars (10–20 μm thick), as the grid bars were clearly visible when imaged in SEM. These sample preparations substantially reduced the FIB-milling times while maintaining the random orientations of the crystals, as illustrated by the improvement of reciprocal space sampling (Figure 2).

**Figure 2.**
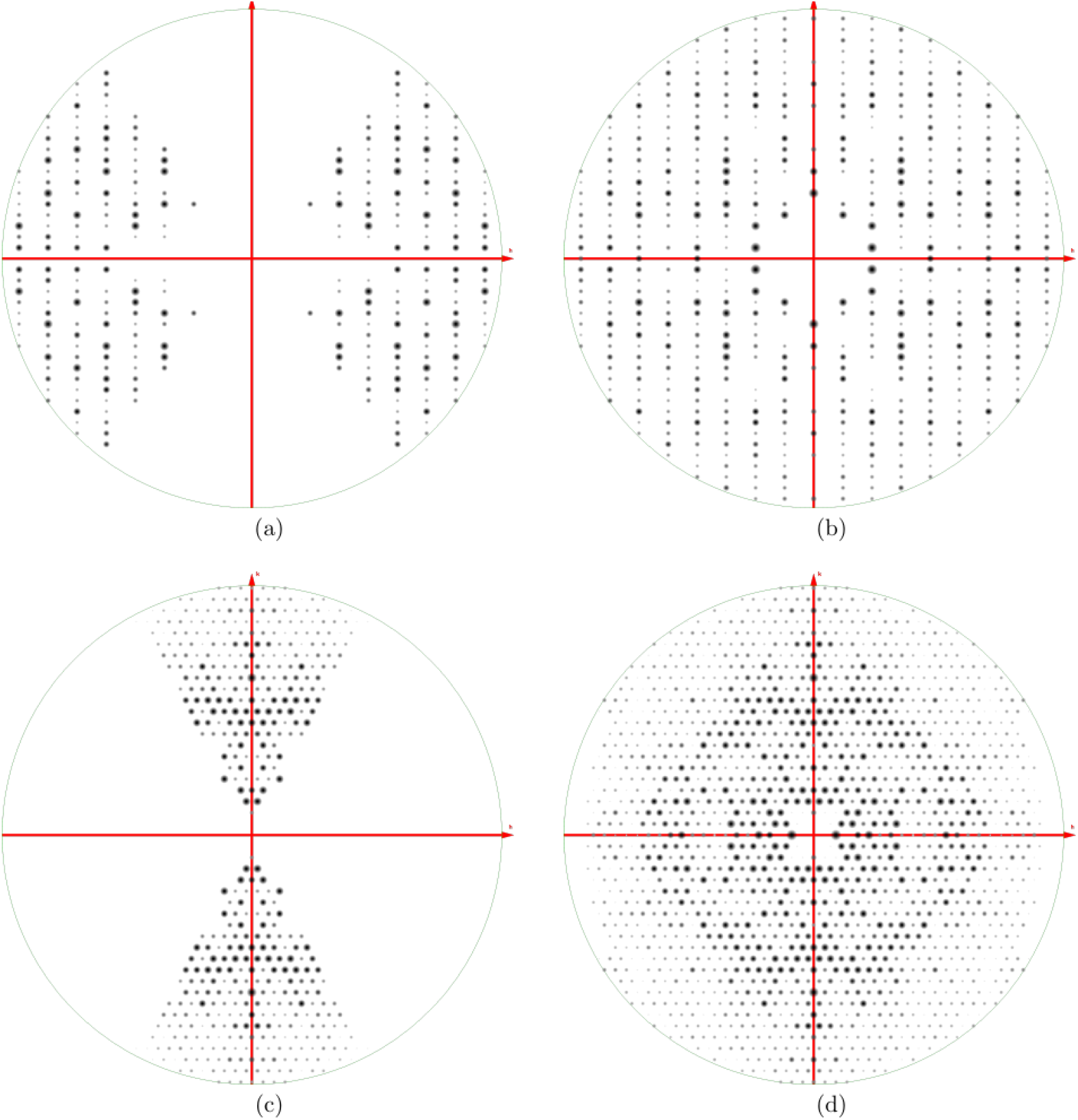
Recovery of missing reflections in MicroED datasets of catalase and Mpro. (a-b) Catalase. (a) The *h*0*l* zone of observed reflections to 8 Å resolution in the preferred orientation dataset viewed along the *k*-axis. (b) The same zone as in (a) for the missing cone eliminated dataset. (c-d) Mpro. (c) The *hk*0 zone of observed reflections to 2.5 Å resolution in the preferred orientation dataset viewed along the *l*-axis. (d) The same zone as in (c) for the missing cone eliminated dataset viewed along the *l*-axis. All plots were rendered with *ViewHKL* from the CCP4 suite.

For the catalase suspended crystal sample, a cryogenically cooled Helios pFIB/SEM (Thermo-Fisher) was used for lamellae milling. Specifically targeting the crystals using an integrated fluorescence microscope (iFLM) and correlative light electron microscopy (CLEM), as in suspended drop crystallization (Gillman *et al*., 2023), was unnecessary because the catalase crystals were at a high concentration in the stock, and capturing a randomly oriented crystal within the lamella was straightforward. During all milling steps, the lamella was imaged using the iFLM set to the 385 nm LED to ensure the presence of randomly oriented crystals. Crystals that appeared as thin rectangles when imaging normal to the grid bars were taken to be in an ideal orientation for capturing the missing cone of data. For the Mpro suspended crystal sample, a cryogenically cooled Aquilos FIB/SEM (Thermo-Fisher) was used for lamellae milling. Because the suspended drop of Mpro crystals was briefly blotted, only a minimal volume of sample was retained by the grid bars, which allowed for crystal morphological features to be observed and targeted. In contrast to the instrumental setup used for catalase, the Aquilos used for the Mpro sample does not have the ability to image the samples using fluorescence. We chose to evaluate our sample using this instrument since it is more generally accessible. To capture as many crystals as possible, several long lamellae expanding over the whole grid square were milled, generally containing 2–5 crystals each.

### 3.2. Eliminating the missing cone in the MicroED structures of catalase and Mpro

Catalase was the second protein determined by MicroED (Nannenga, Shi, Hattne *et al*., 2014). A single crystal yielded relatively high completeness, however, even merging data from five additional crystals could not eliminate the missing cone. This is because catalase forms flat rectangular plate-like crystals that, when dispensed on the grid, align with the *c*\*-axis of their lattices normal to the plane of the grid. MicroED data were collected from a suspended crystal of catalase and merged with the previously published data showing preferred orientation (Nannenga, Shi, Hattne *et al*., 2014). Similarly, Mpro yields flat plate-like crystals with preferred orientations on the grids. In addition, the crystal symmetry of the Mpro crystals is low, leading to low completeness of our initially collected supported crystals (∼60%). These preferred orientation data were combined with Mpro data collected of the sample in suspension. We produced two separate datasets for each protein: one “preferred orientation” dataset and one “missing cone eliminated”. These datasets were processed and refined separately, and the observed reflections, statistics, and refined maps were subsequently compared. Data processing statistics for both “preferred orientation” and “missing cone eliminated” catalase and Mpro datasets are presented in Table 1.

**Table 1.**
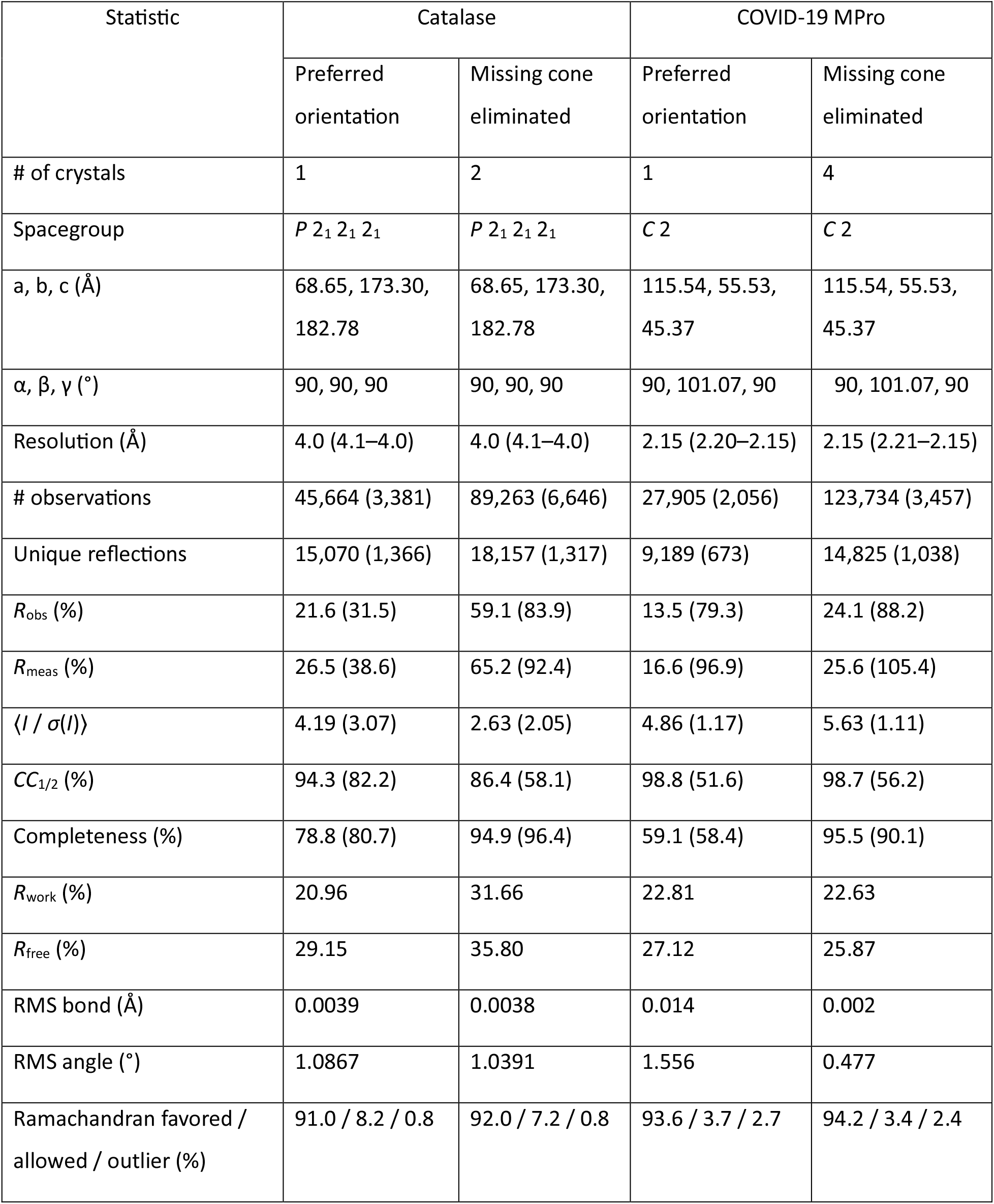
Preferred orientation vs. missing cone eliminated dataset statistics for catalase and Mpro.

The zone at *k* = 0 clearly illustrates that a significant amount of the missing cone of reciprocal space was recovered (Figure 2a, b) for catalase, consistent with the total number of unique reflections increasing from 15,070 to 18,157 (Table 1) within 4 Å resolution. For Mpro, viewing 2D slices along the *l*-axis, the missing cone of data was also clearly recovered (Figure 2c, d), increasing the number of unique reflections from 9,189 to 14,825 (Table 1). As expected, the completeness across the resolution shells of catalase improved after merging the missing cone dataset with the preferred orientation dataset (Figure 3a), increasing the overall dataset completeness from 78.8 to 94.9% (Table 1). Similarly, the completeness for Mpro significantly improved (Figure 3c) from 59.1 to 95.5% (Table 1).

**Figure 3.**
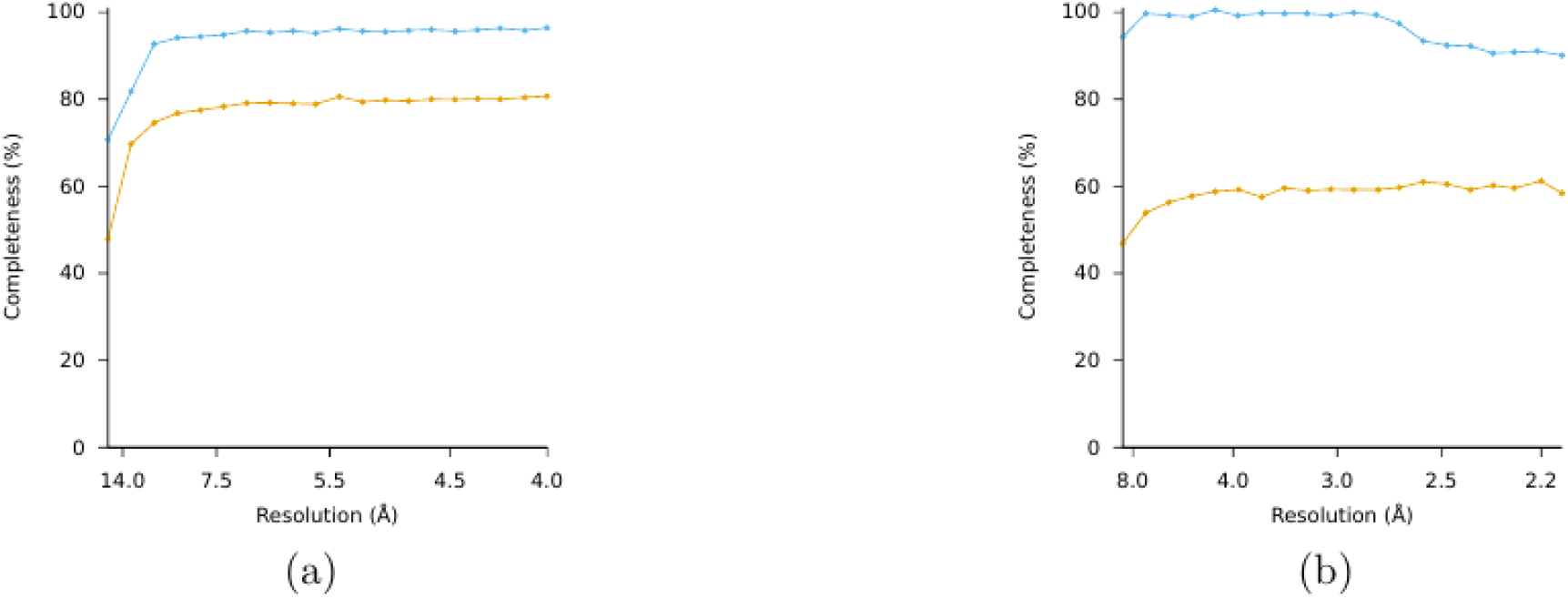
Completeness for preferred orientation data vs. missing cone eliminated data for catalase and Mpro. (a) Completeness (%) of catalase, (b) Completeness (%) of Mpro as functions of resolution (Å). Preferred orientation data in orange, missing cone eliminated data in blue.

Finally, the density maps resulting from the preferred orientation datasets and the missing cone eliminated datasets were compared for each protein (Figure 4). Once the missing cone is eliminated, many regions that were originally poorly resolved show contiguous densities that are easily interpretable. For each protein sample, several instances are illustrated wherein the improved maps facilitated more precise positioning of amino acid sidechains during the construction of the molecular models.

**Figure 4.**
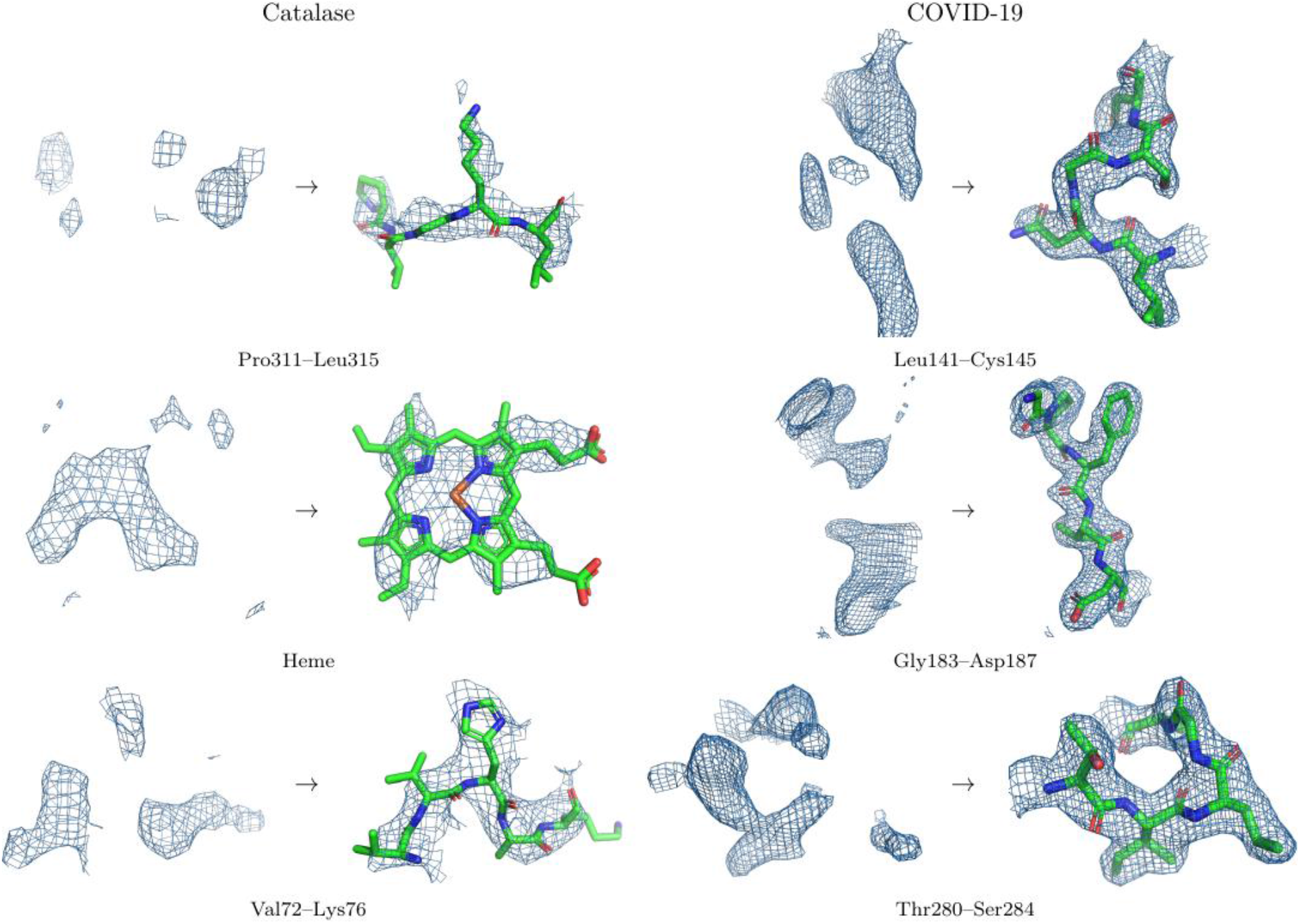
Density improvements upon completion of the reciprocal space. Several regions that exhibit significant density improvement are presented. 2*mF*_*o*_-*DF*_*c*_ maps are all contoured at 1.2 σ above the mean and carved at 2 Å around the model. On the left in each panel, the preferred orientation map density is compared to that for the missing cone eliminated on the right.

## 4. Conclusions

Here, we report a method that enables direct targeting and capturing of the critical missing cone of MicroED data for crystals that systematically adopt a preferred orientation on the EM grid. Suspended crystals of both bovine liver catalase and the COVID-19 main protease Mpro were used as test samples. In both cases, crystals grew as rectangular plates that would normally lie flat on the carbon support film of the grid, such that their crystallographic axes would preferentially orient with the normal of the grid. However, the absence of a support film precluded these crystal plates from having a surface to align with and were effectively frozen in random orientations as postulated in earlier studies (Gillman et al. 2023). Whereas typically no amount of data collection and merging can recover the missing cone due to preferred orientation, crystals of catalase and Mpro were here suspended in essentially random orientations by removing the support film. Crystals are left exposed for FIB-milling and MicroED collection of complete datasets, which yield densities that are interpretable and further facilitate modeling and refinement.

Crystals with a systematic preferential orientation are not common, and we estimate that less than 10% of all protein crystals would fall under this category. The majority of samples will likely adopt random orientations on the grid and may also have higher crystal symmetry. In such cases, even a single nanocrystal can be sufficient for a complete structure by MicroED. However, the missing cone creates a substantial bottleneck in structure solution for crystals that adopt a preferred orientation, and this phenomenon might be more prevalent for smaller crystal sizes. While the suspended drop crystallization approach described here eliminated the preferential orientation problem for MicroED, the missing cone problem remains a prevalent issue in cryotomography (Barth *et al*., 1988). Even for single particle reconstructions, the sample may preferentially orient at the water-air interface (Glaeser & Han, 2017) and innovative approaches may be needed to eliminate the preferential orientation problem.

As the methodologies for MicroED sample preparation are improved and continue to evolve, we anticipate the method to deliver structures for samples that remain beyond the technological reach of other structural biology methods. The innovative solution presented here proves indispensable for crystals exhibiting preferred orientations, unlocking new possibilities for structure determination in MicroED workflows.

## Data availability

Coordinates and maps were deposited in the protein data bank and the EM Data bank (XXXX and YYYY).

## Acknowledgments

The authors would like to thank Dr. Steve Halaby for early work on Mpro. This study was supported by the National Institutes of Health P41GM136508 and the Department of Defense HDTRA1-21-1-0004. The Gonen laboratory is supported by funds from the Howard Hughes Medical Institute. E.D. thanks the Wenner-Gren Foundations for their support through the Wenner-Gren Postdoctoral Fellowship.

## Notes

### Competing Interest Statement

The authors have declared no competing interest.

### Summary of Updates

Update Table 1.

## References

Afonine, P. V., Grosse-Kunstleve, R. W., Echols, N., Headd, J. J., Moriarty, N. W., Mustyakimov, M., Terwilliger, T. C., Urzhumtsev, A., Zwart, P. H. & Adams, P. D. (2012). Acta Crystallogr D Biol Crystallogr 68, 352–367.

Andi, B., Kumaran, D., Kreitler, D. F., Soares, A. S., Keereetaweep, J., Jakoncic, J., Lazo, E. O., Shi, W., Fuchs, M. R., Sweet, R. M., Shanklin, J., Adams, P. D., Schmidt, J. G., Head, M. S. & McSweeney, S. (2022). Sci Rep 12, 10.1038/S41598-022-15930-Z.

Barth, M., Bryan, R. K., Hegerl, R. & Baumeister, W. (1988). Scanning Microsc Suppl 2, 277–284.

Bruhn, J. F., Scapin, G., Cheng, A., Mercado, B. Q., Waterman, D. G., Ganesh, T., Dallakyan, S., Read, B. N., Nieusma, T., Lucier, K. W., Mayer, M. L., Chiang, N. J., Poweleit, N., McGilvray, P. T., Wilson, T. S., Mashore, M., Hennessy, C., Thomson, S., Wang, B., Potter, C. S. & Carragher, B. (2021). Front Mol Biosci 8, 648603.

Clabbers, M. T. B., Martynowycz, M. W., Hattne, J. & Gonen, T. (2022). J Struct Biol X 6, 100078.

Danelius, E., Porter, N. J., Unge, J., Arnold F. H., Gonen T. (2023). J Am Chem Soc 145, 7159–7165.

Emsley, P., Lohkamp, B., Scott, W. G. & Cowtan, K. (2010). Acta Crystallogr D Biol Crystallogr 66, 486–501.

Foroughi, L. M., Kang, Y. N. & Matzger, A. J. (2011). Cryst Growth Des 11, 1294–1298.

Gallagher-Jones, M., Glynn, C., Boyer, D. R., Martynowycz, M. W., Hernandez, E., Miao, J., Zee, C.-T., Novikova, I. V, Goldschmidt, L., McFarlane, H. T., Helguera, G. F., Evans, J. E., Sawaya, M. R., Cascio, D., Eisenberg, D. S., Gonen, T. & Rodriguez, J. A. (2018). Nat Struct Mol Biol 25, 131–134.

Gillman, C., Nicolas, W. J., Martynowycz, M. W. & Gonen, T. (2023). IUCrJ 10, 10.1107/s2052252523004141.

Glaeser, R. M. & Downing, K. H. (1993). Ultramicroscopy 52, 478–486.

Glaeser, R. M. & Han, B.-G. (2017). Biophys Rep 3, 1–7.

Griner, S. L., Seidler, P., Bowler, J., Murray, K. A., Yang, T. P., Sahay, S., Sawaya, M. R., Cascio, D., Rodriguez, J. A., Philipp, S., Sosna, J., Glabe, C. G., Gonen, T. & Eisenberg, D. S. (2019). Elife 8, e46924.

Hattne, J., Reyes, F. E., Nannenga, B. L., Shi, D., Cruz, M. J. de la, Leslie, A. G. W. & Gonen, T. (2015). Acta Crystallogr A Found Adv 71, 353–360.

Hattne, J., Shi, D., Glynn, C., Zee, C.-T., Gallagher-Jones, M., Martynowycz, M. W., Rodriguez, J. A. & Gonen, T. (2018). Structure 26, 759--766.e4.

Hattne, J., Clabbers, M. T. B., Martynowycz, M. W., Gonen, T. (2023). Structure 31, 1504–1509.

Kabsch, W. (2010a). Acta Crystallogr D Biol Crystallogr 66, 133–144.

Kabsch, W. (2010b). Acta Crystallogr D Biol Crystallogr 66, 125–132.

Martynowycz, M. W., Clabbers, M. T. B., Hattne, J. & Gonen, T. (2022). Nat Methods 19, 724–729.

Martynowycz, M. W., Clabbers, M. T. B., Unge, J., Hattne, J. & Gonen, T. (2021). Proc Natl Acad Sci U S A 118, 1–7.

Martynowycz, M. W. & Gonen, T. (2021). STAR Protoc 2, 100686.

Martynowycz, M. W., Shiriaeva, A., Clabbers, M. T. B., Nicolas, W. J., Weaver, S. J., Hattne, J. & Gonen, T. (2023). Nat Commun 14, 1086.

Martynowycz, M. W., Zhao, W., Hattne, J., Jensen, G. J. & Gonen, T. (2019a). Structure 27, 545-548.e2.

Martynowycz, M. W., Zhao, W., Hattne, J., Jensen, G. J. & Gonen, T. (2019b). Structure 27, 1594-1600.e2.

Mastronarde, D. N. (2003). Microscopy and Microanalysis 9, 1182–1183.

Matricardi, V. R., Moretz, R. C. & Parsons, D. F. (1972). Science (1979) 177, 268–270.

McCoy, A. J., Grosse-Kunstleve, R. W., Adams, P. D., Winn, M. D., Storoni, L. C. & Read, R. J. (2007). J Appl Crystallogr 40, 658–674.

Mu, X., Gillman, C., Nguyen, C. & Gonen, T. (2021). Annu Rev Biochem 90, 431–450.

Nannenga, B. L., Shi, D., Hattne, J., Reyes, F. E. & Gonen, T. (2014). Elife 3, e03600.

Nannenga, B. L., Shi, D., Leslie, A. G. W. & Gonen, T. (2014). Nat Methods 11, 927–930.

Purdy, M. D., Shi, D., Chrustowicz, J., Hattne, J., Gonen, T. & Yeager, M. (2018). Proc Natl Acad Sci U S A 115, 13258–13263.

Shi, D., Nannenga, B. L., Iadanza, M. G. & Gonen, T. (2013). Elife 2, e01345.

Shiriaeva, A., Martynowycz, M. W., Nicolas, W. J., Cherezov, V. & Gonen, T. (2023). Biorxiv: The Preprint Server.

Stahlberg, H., Biyani, N. & Engel, A. (2015). Arch Biochem Biophys 581, 68–77.

Sumner, J. B. & Dounce, A. L. (1937). Science (1979) 85, 366–367.

Yonekura, K., Kato, K., Ogasawara, M., Tomita, M. & Toyoshima, C. (2015). Proceedings of the National Academy of Sciences 112, 3368–3373.

